# The effects of biodegradable mulch film on the growth, yield, and water use efficiency of cotton and maize in an arid region

**DOI:** 10.1101/703264

**Authors:** Lu Deng, Ruide Yu, Qian Wang

## Abstract

Plastic residual film pollution in China is serious, and the use of degradable mulch film instead of plastic mulch can effectively alleviate this situation. The substitution of common polyethylene plastic mulch film with biodegradable mulch film in the agricultural production of cotton and maize in an arid region was investigated in the present study. Using bare soil as the control, we compared the effects of common polyethylene plastic film and biodegradable mulch film on crop growth, yield, and water use efficiency (WUE) in maize and cotton. The results indicated that: (1) the biodegradable mulch film in this region remained intact for 60 days after being laid down, significantly degrading after 120 days, and was associated with increased soil temperature, moisture conservation, and degradability in comparison to a bare soil control. (2) Both the biodegradable mulch film and the polyethylene plastic film significantly increased various physiological parameters, such as crop height, stalk diameter, and leaf area. (3) The biodegradable mulch film significantly increased maize and cotton crop yield by 69.4–76.2% and 65.2–71.9%, respectively, compared to the bare soil control. (4) Compared to the bare soil control, the biodegradable mulch film effectively increased WUE in the crops by 64.5–73.1%. In summary, biodegradable mulch film had comparable results to the common polyethylene plastic film in increasing soil temperature, moisture conservation, crop growth, yield, and WUE. As the biodegradable mulch film causes no residual pollution, it is thus preferable to common plastic mulch film for agricultural applications in arid regions and supports the sustainable development of agroecosystems.

## 1. INTRODUCTION

With a total land area of 166 million hm^2^, of which 63,084,800 hm^2^ constitutes agricultural regions, Xinjiang contributes significantly to agricultural security in China[1]. Xinjiang has an average precipitation and evaporation capacity of 154.5 mm and 1,600–2,200 mm^2^, respectively, which results in water shortages that limit agricultural development in this region[2]. According to the National Bureau of Statistics of China (http://data.stats.gov.cn/index.htm), the total agricultural water consumption in Xinjiang was 93.1% of the total water supply in 2017. Due to this high agricultural water consumption, it is necessary that agricultural water usage efficiency is increased in order to support ecological conservation and industrial development, which are important for local economic and social development.

According to data released by the National Bureau of Statistics of China, the use of plastic film for agricultural use in China has increased from 1.845 million tons in 2006 to 2.6 million tons in 2015.(data from the National Bureau of Statistics of China, http://www.stats.gov.cn/).Xinjiang is a typical arid region where plastic film is widely used in agricultural production[3] for increasing soil temperature and moisture conservation, as well as for increasing crop yield[4–8]. Polyethylene plastic degrades poorly and leaves residual film for a long period of time, causing severe pollution[9–11]. Xinjiang is one of the major regions in China experiencing major residual film pollution. The area has a residual film content of 262– 597 kg/hm^2^, which has seriously impacted on the sustainable development of local agriculture. Using environmentally-friendly, controllable biodegradable mulch film to replace common plastic film can increase crop yield, reduce agricultural water consumption, and effectively resolve the issues related to residual film pollution[3]. It is one of the technologies that possess the most potential for promoting the sustainable development of agriculture. Han et al.[12], Chen et al.[13] and Ren et al.[4,5] found that in the Loess Plateau, China, biodegradable mulch film could improve maize yield and water use efficiency (WUE) compared to no film. Moreno et al.[14] found that biodegradable mulch film increased tomato yield in comparison to no film in Central Spain. Yao et al.[15] showed that in Hubei Province, biodegradable mulch film could improve rice production and reduce the emissions of the greenhouse gases CH_4_ and N_2_O. However, the above-mentioned studies have some shortcomings. For instance, the effectiveness of biodegradable mulch film is closely correlated with environmental conditions (temperature, precipitation, etc.) and management practices. The degree of degradation and degradation period have been found to differ in different areas, and thus the ultimate effectiveness of biodegradable mulch film varies[3]. Most studies on biodegradable mulch film have focused on semi-arid regions where the annual precipitation and temperature vary from 435–550 mm and 3.6–14.3°C, respectively. However, arid regions experience more extreme environmental conditions, including major differences in day and night temperature (daily mean temperature between −25°C to 30°C), low precipitation (annual precipitation < 200 mm), and strong solar radiation. Very few studies have evaluated the efficacy of biodegradable mulch film in arid regions. Additionally, most of the studies mentioned above only focused on the final crop yield, rather than tracking the entire crop growth process. These studies also only focused on the impact of biodegradable mulch film on a single crop and thus lack a comparative assessment of the effects in different crops. Finally, these studies only assessed the effects of biodegradable mulch film on crop yield and the external environment, rather than focusing on the degradation of the biodegradable mulch film itself.

To address the inadequacies mentioned above, and to further provide a theoretical basis and technical support for the application and promotion of biodegradable mulch film in agricultural practices, the present study comparatively analyzed the differences in the effects of biodegradable mulch film and common polyethylene plastic film on the growth characteristics, yield increase, and WUE of maize and cotton in an arid region in Northwestern China. The major aims of this study were to: (1) investigate whether the positive impacts of biodegradable mulch film in the arid region on crop physiological characteristics could match those of common plastic film; (2) investigate whether biodegradable mulch film in the arid region could significantly increase crop biomass and yield, and whether this differed from common plastic film; (3) comparatively analyze the effect of biodegradable mulch film and common plastic film on WUE; (4) comparatively analyze the degradation of biodegradable mulch film and common plastic film.

## 2. MATERIALS AND METHODS

### 2.1. Site description

The experimental site was located in Shangsanqi Village in Erliugong Town, Changji City, Xinjiang Uygur Autonomous Region, China (87°13′E, 44°02′N; altitude 574 m) between 2015 and 2017. The location is shown in detail in Fig. 1.

**Fig. 1.**
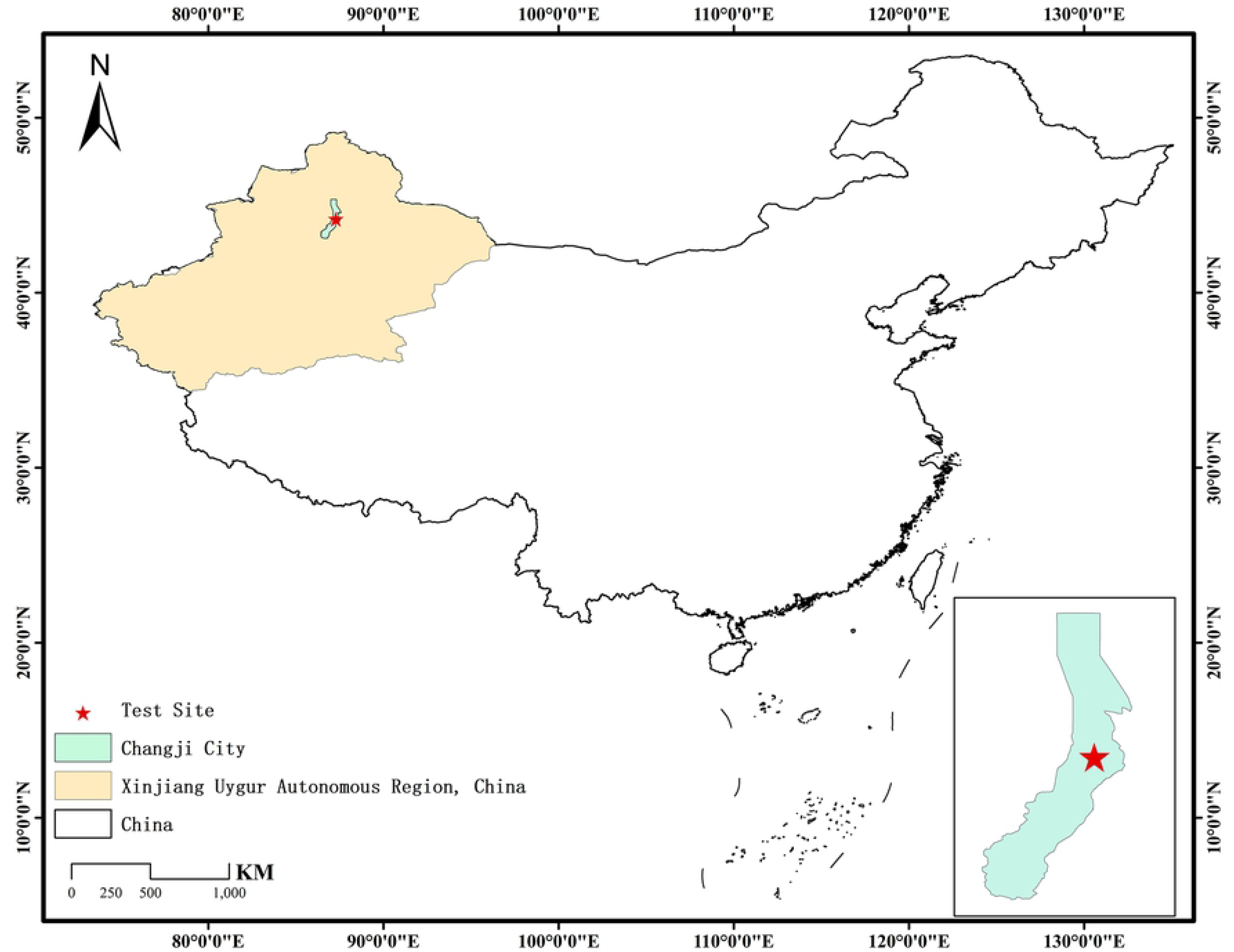
Schematic diagram of the experimental site.

The experimental site has a typical continental temperate arid climate, with cold winters and hot summers, as well as large variations in day and night temperatures. The soil mellowing level is relatively poor in the experimental site, with low fertility levels. At the 0–60 cm soil layer, the soil organic matter content is 3.07–9.68 g•kg^−1^ and the total nitrogen content is 0.15–0.30 g⋅kg^−1^. From 2015 to 2017, the annual precipitation was 174.5 mm, 187.6 mm, and 97.5 mm, and the mean temperature was 7.8°C, 7.1°C, and 7.6°C. Precipitation and temperature at the experimental site are shown in Fig. 2. Low temperatures during early growth stages and lack of sufficient precipitation during the entire growth period are the major factors that limit crop growth in this region.

**Fig. 2.**
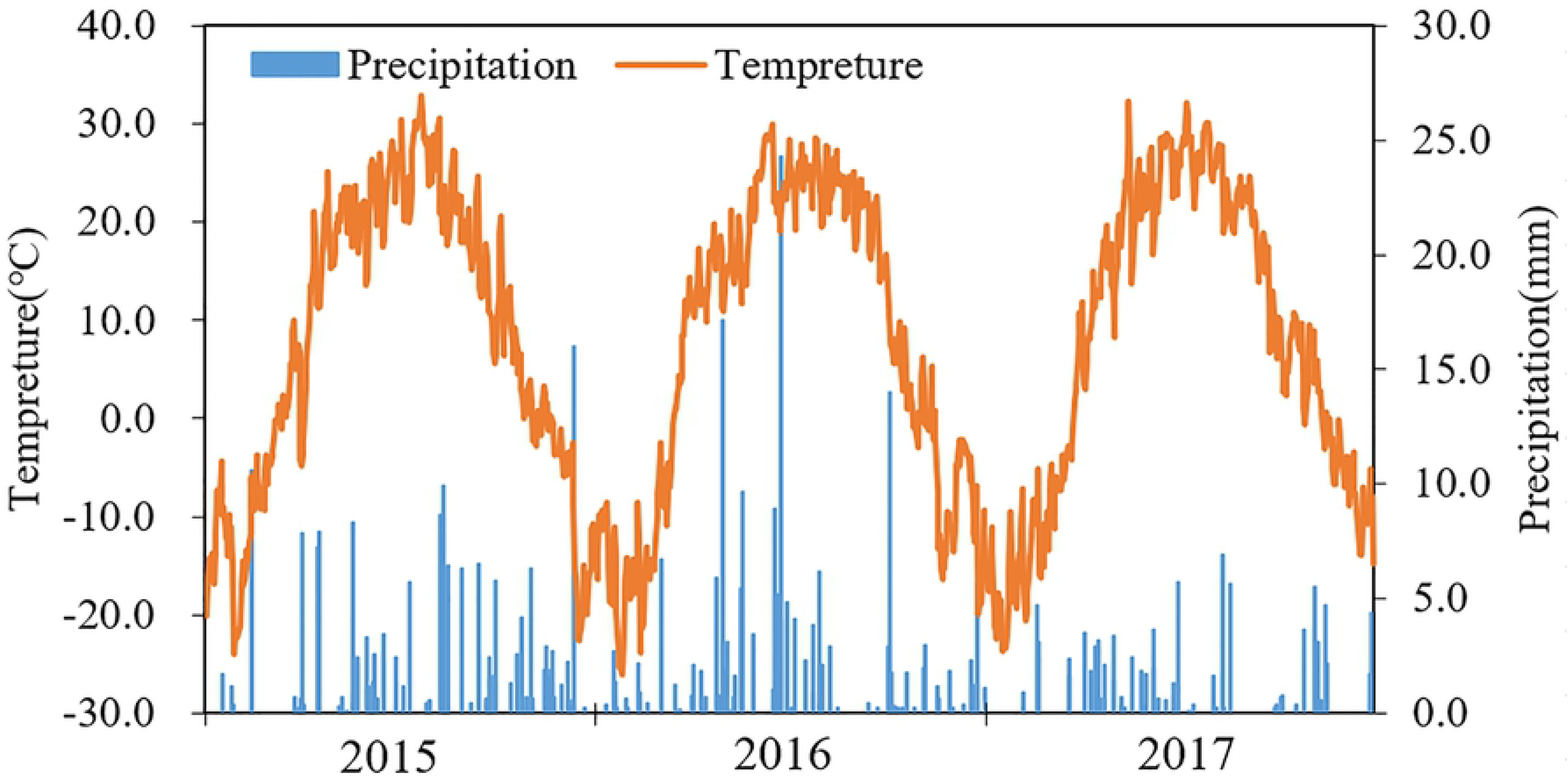
Changes in daily temperature and precipitation at the experimental site from 2015 to 2017.

### 2.2 Field management and experimental design

#### 2.2.1 Experimental materials

Xinjiang’s main crops - maize and cotton, were selected as research objects. The mulch films were obtained from Xinjiang Blue Ridge Tunhe Chemical Industry Joint Stock Co., Ltd. (Xinjiang, China). The experiment was set up as four groups: common plastic film, black biodegradable mulch film, clear biodegradable mulch film, and no mulch. The treatments were as follows: A, clear common plastic film predominantly composed of polyethylene, 0.010 mm thickness. B, clear mulch film predominantly composed of polybutyrate (PBAT ≥ 95%), 0.010 mm thickness. C, black mulch film predominantly composed of (PBAT) (≥ 95%), 0,010 mm thickness. D, unmulched.

#### 2.2.2 Management measures

For maize and cotton, the four treatments (A, B, C, and D) mentioned above were established in the experimental site. Each treatment plot was 30 m in length and 20 m in width. This study was a completely randomized controlled study with six replicates for each treatment. The plots were randomly arranged. The experiments lasted for three years, and the sowing and harvesting dates for each year are shown in Table 1.

**Table 1.** Sowing and harvesting date of the maize and cotton in each year during the experiment.

| Year |  | Sowing date | Harvesting date |
| --- | --- | --- | --- |
| Maize | 2015 | April 27 | September 15 |
|  | 2016 | April 25 | September 16 |
|  | 2017 | April 27 | September 15 |
| Cotton | 2015 | April 20 | October 12 |
|  | 2016 | April 18 | October 23 |
|  | 2017 | April 21 | October 25 |

For the maize treatment, 35 kg P ha^−1^ diammonium phosphate was applied to the experimental plots as the base fertilizer. The films were laid out and seeds were sown using an integrative machine. Maize was sown with a row interval of 60 cm, with 25 cm spacing between each plant. From around April 27 to August 17, 45 mm of drip irrigation was applied every 10 days. From the second to the fifth irrigation, 50 kg N ha^−1^ urea was added to the water. During the growth period, the field management measures remained the same in the four treatments.

For the cotton treatment, 80 kg P ha^−1^ diammonium phosphate was applied to the experimental plots as the base fertilizer. The films were laid out and seeds were sown using an integrative machine. Cotton seeds were sown with a row interval of 60 cm, with 10 cm spacing between each plant. From around April 20 to September 1, water was supplied every 15 d (10 times total). In the early and later stages, 25 mm water was applied; in the middle stage, 50 mm of water was applied. From April 20 to July 30, 80 kg N ha^−1^ urea was applied with the irrigation a total of four times. During the growth period, the field management measures remained the same for all four treatments.

#### 2.2.3 Parameters tested and methods

In the middle of each growth stage, the growth and development of the maize and cotton in the different treatments were observed and recorded, and the emergence rate was calculated. The plant height, stalk diameter, and leaf area of the crops under the different treatments were measured. In addition, at the boll-opening stage, the boll number per plant and boll weight were recorded. Furthermore, the aboveground and underground parts of the maize and cotton plants in six areas of 1 m^2^ were randomly collected. After removing impurities such as soil and sand, the samples were chopped, dried in an oven at 105 ° C for 2 hours, then dried at 80 ° C to constant weight, and weighed to obtain the total biomass in 1 m^2^, as well as the weight of the maize kernel and cotton. From this, we calculated the biomass and yield in 1 hectare.

The degradation of the mulch was graded according to Yang H.D.’s research[16]. The degradation of the mulch film was observed every 10 d after being installed. A 0–5 grade rating was used to define the level of mulch film degradation. Grade 0 represents intact mulch film with no cracks. Grade 1 represents the appearance of the first crack. Grade 2 represents the appearance of small cracks in 25% of the field. Grade 3 represents the appearance of 2– 2.5cm-long cracks. Grade 4 represents the appearance of evenly distributed, network-like cracks. Grade 5 represents the breakage and degradation of the pieces into fragments smaller than 4 cm × 4 cm.

Water use efficiency (kg⋅hm^−2^⋅mm^−1^) = grain yield (kg⋅hm^−2^)/total water consumption (mm). Total water consumption was the sum of precipitation and irrigation water usage.

## 3. RESULTS

### 3.1 Effect of different mulch film treatments on crop physiological characteristics

Compared to the bare soil, the three mulch films greatly affected the emergence rate and growth progression of the maize and cotton (Table 2). The emergence rate of both crops was similar in the three mulch film treatments, all of which were higher than in the bare soil (*P* < 0.05). The emergence rate of the maize and cotton was in the following order: A > B > C > D. The emergence rate of the maize and cotton under the mulch film treatments was 56.1% and 62.4% higher, respectively, than that under the bare soil. For maize, the emergence times in treatment A and B were 11.3 d and 0.7 d earlier than in treatment C, and the emergence time was significantly shorter in the three mulch film treatments than that under bare soil (*P* < 0.05). For cotton, the emergence time under the different treatments was consistent with the pattern observed in maize. Treatment A had the shortest emergence time of 10.8 d, while the B and C treatments had similar emergence times of 12.1 d and 12.8 d, respectively, both of which were significantly earlier than treatment D at 16.5 d (*P* < 0.05). In terms of the time taken from sowing to harvesting the maize, treatment A was shortest at 129.7 d. Treatment B and C took 135.0 d and 143.7 d, respectively. Treatment D took the longest time of 145 d. In terms of the time taken from cotton sowing to boll opening, the difference in time was consistent with that of maize: treatment A took the shortest time of 111.5 d, while treatment B and C took 112.3 d and 113.1 d, respectively. Treatment D took 117.2 d. (4) As plant growth progressed, when comparing treatment A with B and C, the differences in the time required for growth became increasingly significant. Furthermore, the time needed for growth to progress in B and C became increasingly similar to D, particularly in treatment C. It is possible that in the early stages, the biodegradable mulch films were intact and had a similar effect as the common plastic film, but as the biodegradable mulch film gradually degraded, the moisture conservation function decreased, and thus the effect became increasingly similar to that of the bare soil.

**Table 2.**
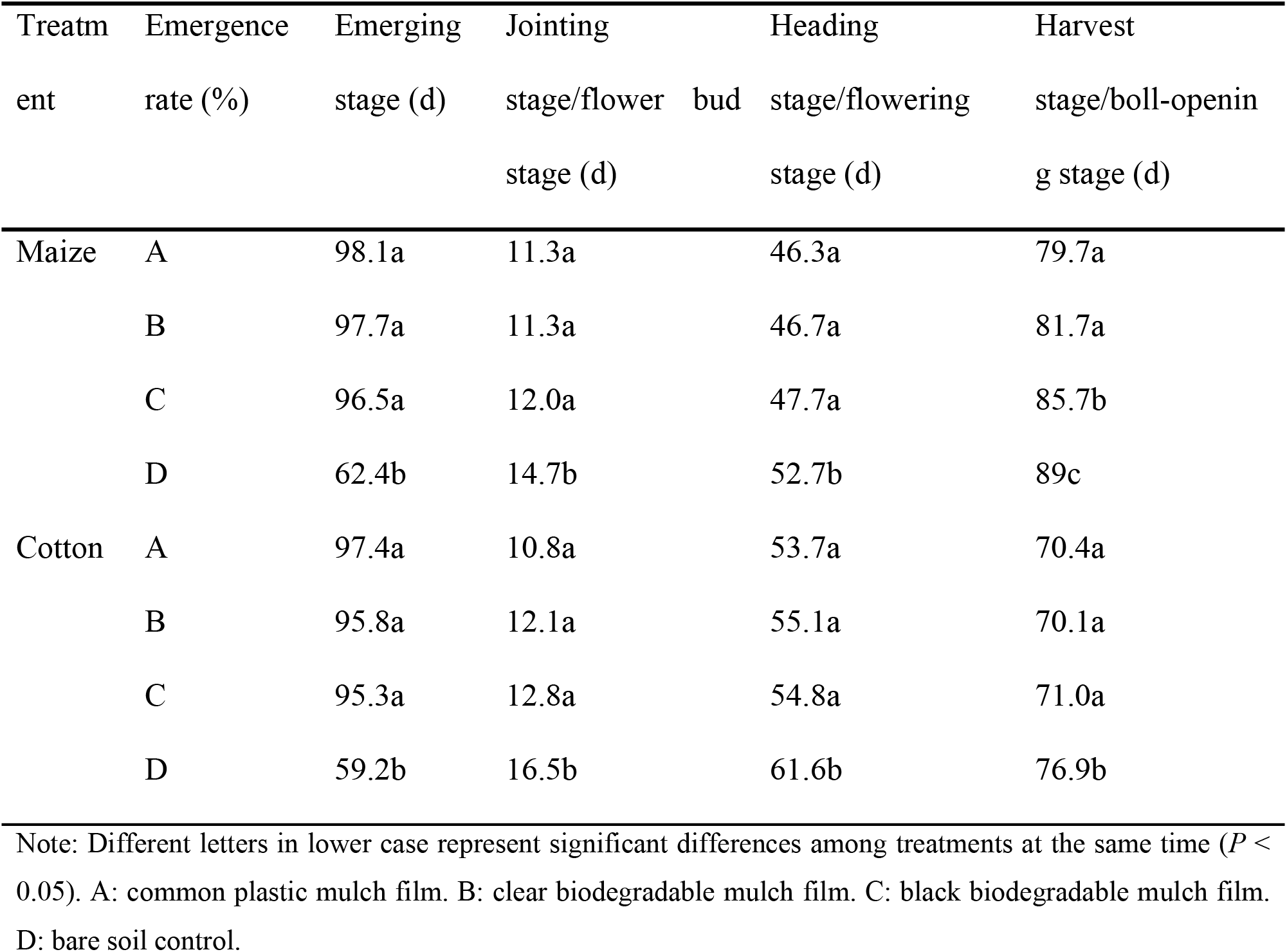
The average emergence rate (%) and growth progression (days after sowing) of maize and cotton under different treatments from 2015 to 2017.

Similarly, compared to the bare soil, the three mulch films also had a large impact on the physiological characteristics of the maize and cotton (Tables 3 and 4). (1) During the entire growth process, treatment A, B, and C significantly increased the plant height, stalk diameter, and leaf area of the maize compared to D (*P* < 0.05). The difference was not significant among treatment A, B, and C. For cotton, the physiological characteristics under the four treatments exhibited a similar pattern as in maize. In each stage, plant height, leaf area, boll number per plant, and boll weight of cotton (boll-opening stage only) did not differ among treatments A, B, and C, all of which were significantly greater than in treatment D (*P* < 0.05).

**Table 3.** The average growth parameters of maize under different treatments from 2015 to 2017.

| Parameter | Treatment | Jointing stage | Heading stage | Harvest stage |
| --- | --- | --- | --- | --- |
| Plant height (cm) | A | 68a | 173a | 188a |
|  | B | 65a | 166a | 189a |
|  | C | 64a | 166a | 191a |
|  | D | 48b | 158b | 175b |
| Stalk diameter (cm) | A | 0.92a | 3.12a | 3.35a |
|  | B | 0.88a | 3.10a | 3.31a |
|  | C | 0.85a | 3.10a | 3.34a |
|  | D | 0.78b | 2.72b | 2.90b |
| Leaf area (m<sup>2</sup> per plant) | A | 0.29a | 0.93a | 0.89a |
|  | B | 0.28a | 0.91a | 0.88a |
|  | C | 0.26a | 0.91a | 0.89a |
|  | D | 0.2b | 0.85b | 0.81b |

Note: Different letters in lower case represent significant differences among treatments at the same time ( $P <$
0.05). A: common plastic mulch film. B: clear biodegradable mulch film. C: black biodegradable mulch film. D: bare soil control.

**Table 4.**
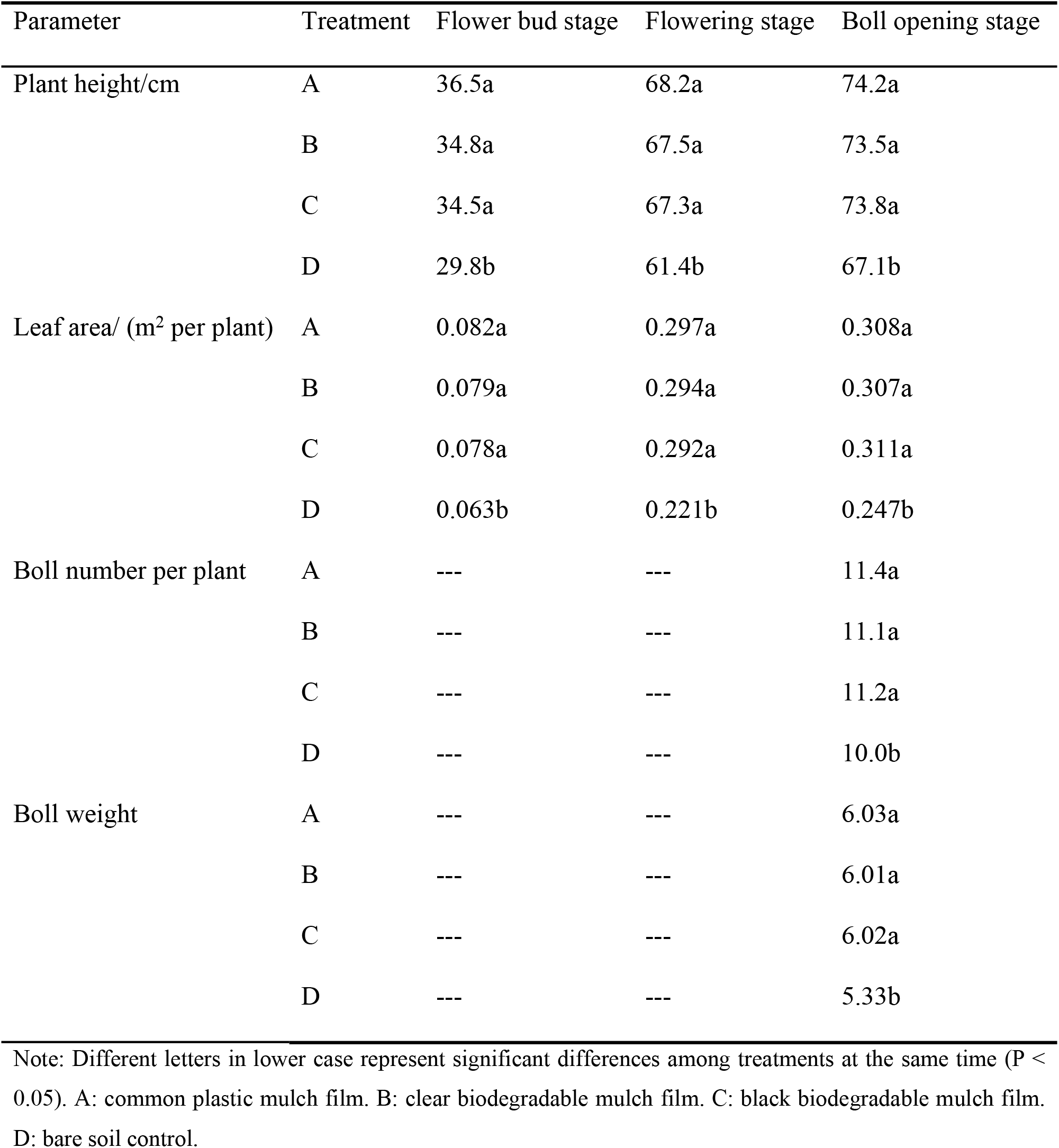
The average growth parameters of cotton under different treatments from 2015 to 2017.

(2) For maize, we found that at the jointing stage, heading stage, and harvest stage, treatment A, B, and C increased plant height by 36.8%, 6.5%, and 8.2%, respectively, compared to treatment D, suggesting that film mulching had the greatest impact on maize plant height at the jointing stage. Treatment A, B, and C increased stalk diameter by 13.2%, 14.2%, and 14.9% at the jointing stage, heading stage, and harvest stage, respectively, compared to treatment D. This indicated that mulch films had a similar effect on maize stalk diameter at each stage. Treatment A, B, and C increased leaf area by 38.3%, 7.8%, and 9.5% at the jointing stage, heading stage, and harvest stage, respectively, compared to D, indicating that film mulching had the most significant impact on maize leaf area during the jointing stage.

(3) For cotton, we found that at the flower bud stage, flowering stage, and boll-opening stage, treatment A, B, and C increased plant height by 18.3%, 10.2%, and 10.0%, respectively, compared to D, suggesting that film mulching affected cotton plant height most significantly at the flower bud stage. At the flower bud stage, flowering stage, and boll-opening stage, treatment A, B, and C increased leaf area by 26.5%, 33.2%, and 25.0%, respectively, compared to treatment D, indicating that film mulching affected cotton leaf area most significantly at the flower bud stage. At the boll-opening stage, treatment A, B, and C on average increased the boll number per plant by 5.0%, and increased boll weight by 12.9%, compared to treatment D, indicating that the mulch film treatments had a significant effect on increasing cotton yield.

(4) For maize, treatment A, B, and C, on average, increased plant height by 17.2% and increased stalk diameter and leaf area by 14.1% and 18.5%, respectively, compared with treatment D. This suggested that the mulch film treatments affected the physiological characteristics of the maize in the following order: leaf area > plant height > stalk diameter. In terms of cotton, treatment A, B, and C, on average, increased plant height by 12.7%, increased leaf area by 28.2%, and increased the boll number per plant by 5.0% and the boll weight by 12.9%, compared to treatment D. This suggested that the mulch film treatments affected the physiological characteristics of the cotton in the following order: leaf area > boll weight > plant height > boll number per plant.

(5) While there were differences among the three mulch films, these differences were not statistically significant. During the jointing stage and heading stage of maize, the plant height, stalk diameter, and leaf area were in the following order: A > B > C (no significant difference, *P* > 0.05). At the mature stage, the plant height, stalk diameter, and leaf area did not differ significantly among the mulch treatments. For cotton, the plant height and leaf area at the flower bud stage and flowering stage did not differ significantly. Similarly, at the boll-opening stage, the plant height, leaf area, boll number per plant, and boll weight did not differ significantly between the mulch treatments.

### 3.2 Effect of different mulch films on crop biomass

The maize biomass under different treatments during the experiment is shown in Fig. 3. The results showed that: (1) in the three years, during the entire growth period, treatment A increased the maize biomass by 27% compared to treatment D; B increased maize biomass by 21% compared to treatment D; and C increased maize biomass by 20.1% compared to treatment D. This indicated that under all three mulch film treatments, maize biomass was significantly higher than the bare soil treatment. Of these treatments, the common plastic film increased the maize biomass most significantly, while the two biodegradable mulch films did not differ significantly from each other.

**Fig. 3.**
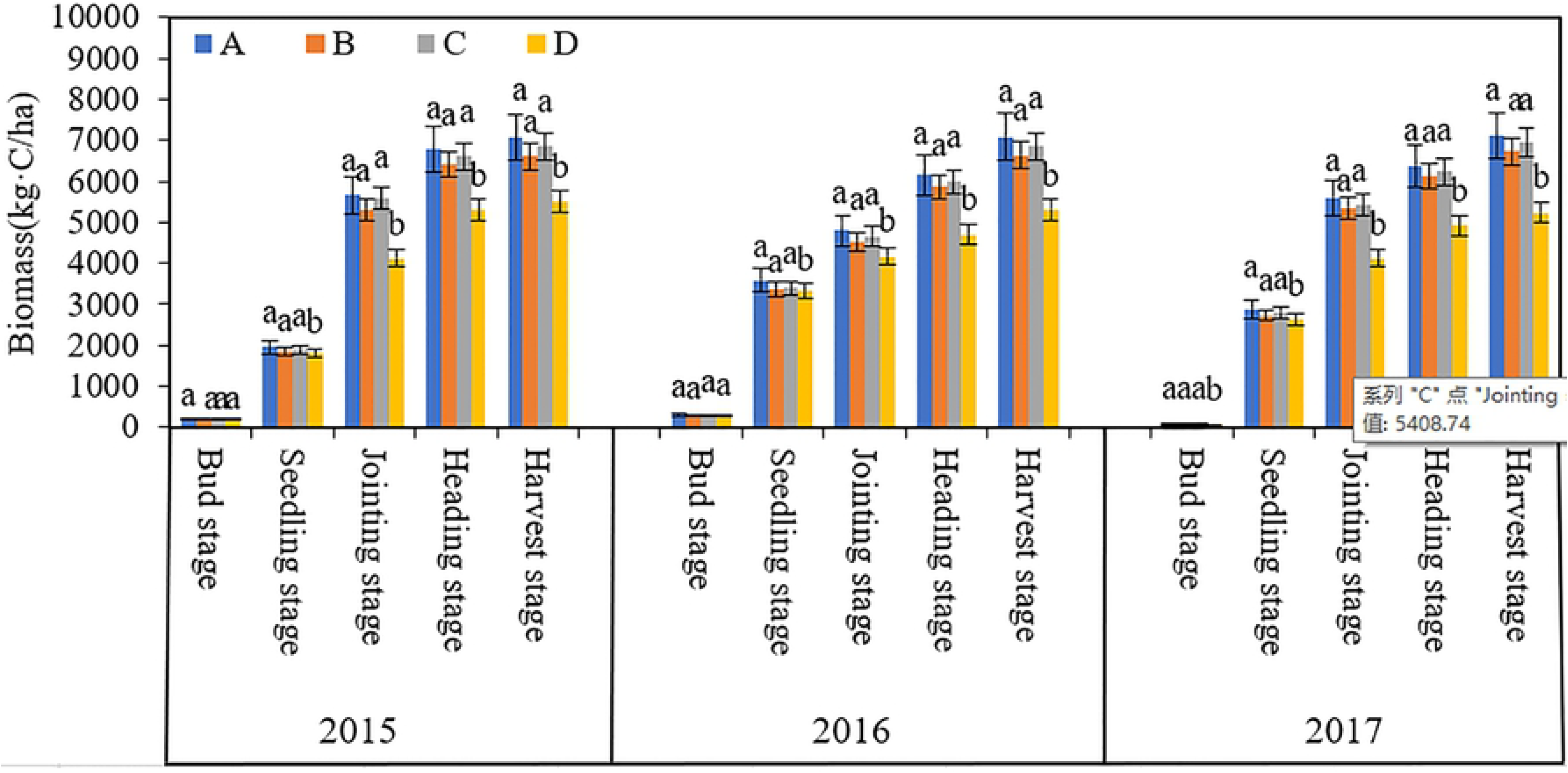
Interannual variations in maize biomass during the growth period under different treatments (2015–2017). The error bars indicate the standard deviation. Different letters in lower case represent significant differences among treatments at the same stage (*P* < 0.05). A: common plastic mulch film. B: clear biodegradable mulch film. C: black biodegradable mulch film. D: bare soil control.

(2) During the three years, treatment A increased the maize biomass by 6.97%, 8.50%, 29.30%, 29.58%, and 32.52% in the bud stage, seedling stage, jointing stage, heading stage, and harvest stage, respectively, compared to treatment D. Treatment B increased the maize biomass by 3.52%, 3.87%, 23.84%, 23.62%, and 25.01% in the bud stage, seedling stage, jointing stage, heading stage, and harvest stage, respectively, compared to treatment D. Compared to treatment D, treatment C increased the maize biomass by −0.10%, 2.39%, 22.58%, 23.40%, and 24.42% in the bud stage, seedling stage, jointing stage, heading stage, and harvest stage, respectively. This showed that the changes in maize biomass during the growth period fitted a logistic shape growth curve, with the film mulching having a greater impact on maize biomass in the later growth stages than in the early growth stages.

Fig. 4 shows cotton biomass under the different treatments during the experiment. In the three treatment years during the entire growth period of cotton, cotton biomass increased by 20.7%, 15.8%, and 12.1% in treatment A, treatment B, and treatment C, respectively, indicating that cotton biomass was significantly higher in all three mulch film treatments compared to the bare soil. Of these treatments, common plastic film was most effective in increasing cotton biomass, while the effects of the two biodegradable mulch films on cotton biomass did not differ significantly from each other.

**Fig. 4.**
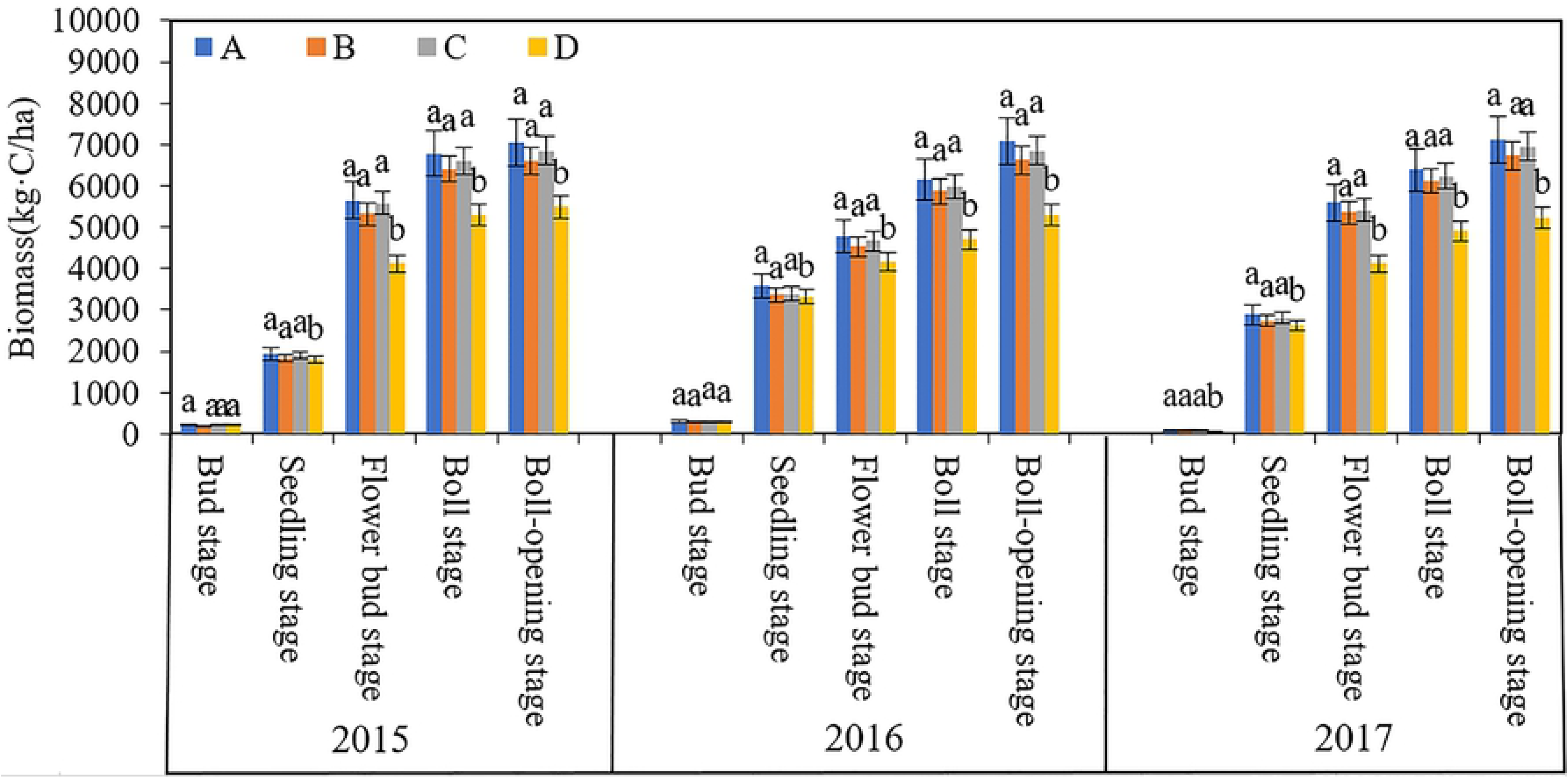
Interannual variations in cotton biomass during the growth period under different treatments (2015–2017). The error bars indicate the standard deviation. Different letters in lower case represent significant differences among treatments at the same stage (*P* < 0.05). A: common plastic mulch film. B: clear biodegradable mulch film. C: black biodegradable mulch film. D: bare soil control.

(2) During the three years, treatment A increased cotton biomass by 16.9%, 12.8%, 11.7%, 30.2%, and 32.1% in the bud stage, seedling stage, flower bud stage, boll stage, and boll-opening stage, respectively, compared to treatment D. Treatment B increased cotton biomass by 9.9%, 7.7%, 7.6%, 25.2%, and 28.6% in the bud stage, seedling stage, flower bud stage, boll stage, and boll-opening stage, respectively, compared to treatment D. In comparison to treatment D, treatment C increased cotton biomass by 2.8%, 4.8%, 4.5%, 22.2%, and 26.0% in the bud stage, seedling stage, flower bud stage, boll stage, and boll-opening stage, respectively. The results showed that the changes in cotton biomass during the growth period fitted a logistic shape growth curve, with the effect of mulch films being greater in the later growth stages than in the early stages.

The interannual variations in cotton biomass showed that the mulch film treatments had a similar effect on biomass accumulation in cotton and maize. Specifically, during the entire growth period, the three mulch film treatments and the unmulched treatment all exhibited a logistic shape growth curve, with biomass being relatively low before the end of May and rapidly increasing thereafter, and then changing slowly after September. All three mulch film treatments significantly increased cotton biomass (*P* < 0.05), but there was no significant difference among the three treatments, although the differences gradually increased during the later growth stages.

The final crop yield during the experiment is shown in Fig. 5. As shown in Fig. 5, the mulch film treatments significantly increased the yield of maize and cotton (*P* < 0.05), but there was no significant difference among the three mulch film treatments (*P* > 0.05). In terms of maize yield, treatment A had the highest average yield of 6,473.6 kg/ha, which was 76.2% higher than in treatment D, and the yield of treatment B was 69.4% higher than treatment D. In treatment C, the yield was 72.6% higher than that in treatment D. Treatment A, B, and C did not differ significantly in average yield over the three years (*P* > 0.05). Of these treatments, treatment A had the highest average yield, followed by treatment C and then treatment B. The variations in cotton yield followed a similar pattern as in maize. Treatment A was associated with the highest cotton yield of 6,575.2 kg/ha, which was 71.9% higher than the yield in treatment D, while the yield of treatment B was 65.2% higher than treatment D. In treatment C, the yield was 69.2% higher than in treatment D. Treatment A, B, and C did not differ significantly in terms of the average cotton yield over the three years (*P* > 0.05). Overall, the different mulch film treatments increased maize yield by 72.7% on average and increased the cotton yield by 68.8% on average. This indicated that the mulch films significantly increased the yield of maize and cotton at the experimental site, and the increase in maize yield was higher than that of cotton.

**Fig. 5.**
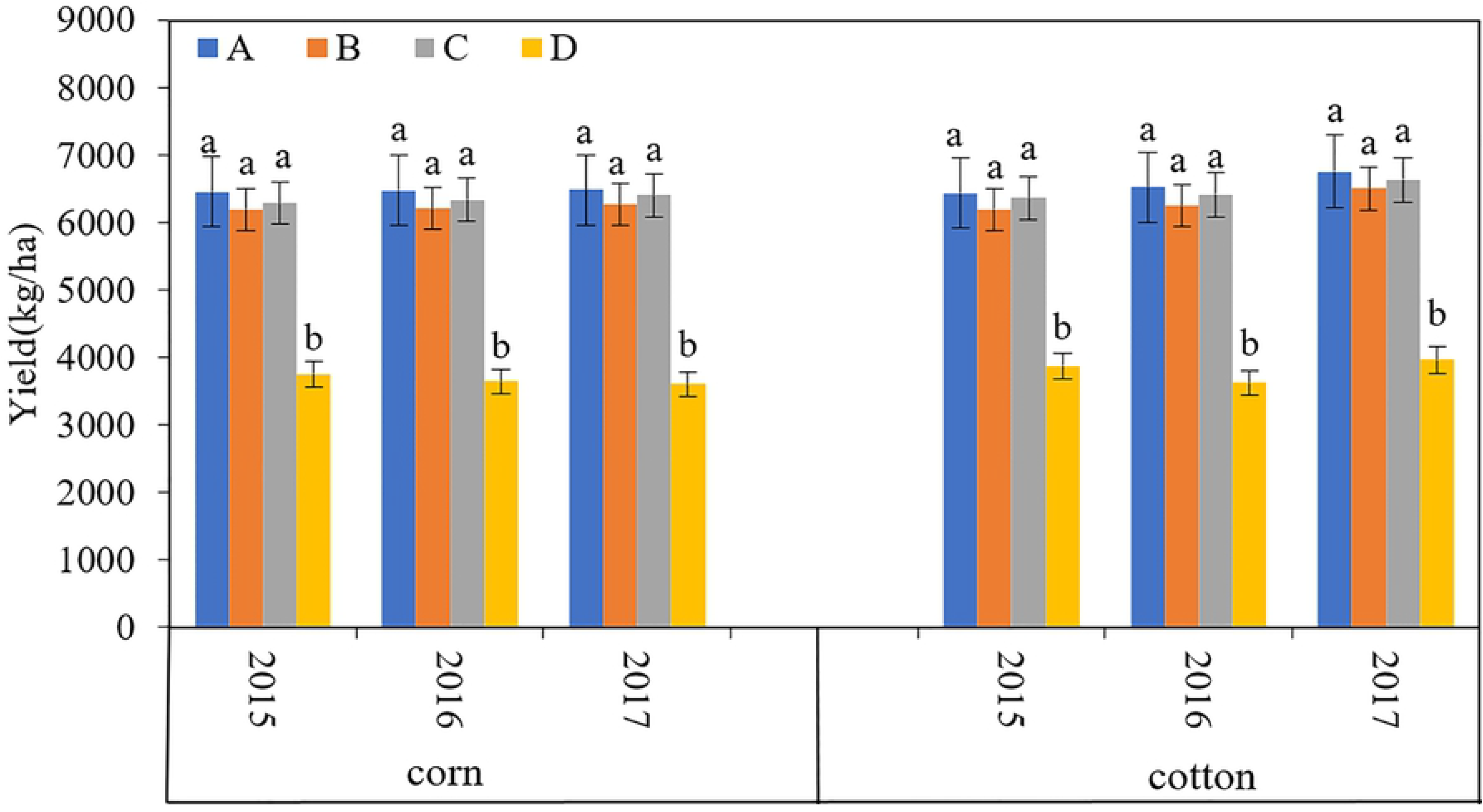
Interannual variations in maize and cotton yield under different treatments (2015– 2017). The error bars indicate the standard deviation. Different letters in lower case represent significant differences among treatments at the same stage (*P* < 0.05). A: common plastic mulch film. B: clear biodegradable mulch film. C: black biodegradable mulch film. D: bare soil control.

### 3.3 WUE

As shown in Table 5, during the three-year experimental period, the mulch film treatments significantly increased the WUE in maize and cotton (*P* < 0.05), but there were no significant differences in the WUE among the three mulch film treatments (*P* > 0.05). In the maize treatment, treatment A had the highest WUE of 10.0 kg⋅hm^−2^/mm, which was 76.2% higher than treatment D. The WUE of treatment B was 69.5% higher than that in treatment D, and treatment C was 73.1% higher than treatment D. The variations in WUE for cotton exhibited a similar pattern as for maize. Treatment A had the highest WUE of 31.9 kg⋅hm^−2^/mm, which was 71.1% higher than treatment D. Treatment B and C had 64.5% and 68.4% higher WUE, respectively, than treatment D. No significant differences in WUE were observed among the mulch treatments during the three years. On average, the three mulch film treatments increased WUE in the maize and cotton by 73.0% and 68.0%, respectively. This indicates that the mulch films significantly increased WUE in maize and cotton at the experimental sites, and the increase in maize was greater than that in cotton.

**Table 5.** Water use efficiency in maize and cotton under different treatments from 2015 to 2017 (kg⋅hm^−2^/mm).

| Treatment |  | A | B | C | D |
| --- | --- | --- | --- | --- | --- |
| Maize | 2015 | 10.0a | 9.6a | 9.8a | 5.8b |
|  | 2016 | 9.2a | 8.8a | 9.0a | 5.2b |
|  | 2017 | 10.7a | 10.4a | 10.6a | 6.0b |
| Cotton | 2015 | 30.6a | 29.4a | 30.2a | 18.4b |
|  | 2016 | 22.6a | 21.7a | 22.2a | 12.6b |
|  | 2017 | 42.4a | 40.8a | 41.7a | 24.9b |

Note: Different letters in lower case represent significant differences among treatments at the same time ( $P < 0.05$ ). A: common plastic mulch film. B: clear biodegradable mulch film. C: black biodegradable mulch film. D: bare soil control.

### 3.4 The degradation rate of the different biodegradable mulch films

The biodegradable mulch films exhibited similar degradation patterns in the cotton and maize treatments. The details are shown in Table 6.

**Table 6.** Degradation rate of the different mulch films used in maize and cotton.

| Year | Treatment | Days after the film was laid down |
| --- | --- | --- |

|  |  |  | 40 | 50 | 60 | 70 | 80 | 90 | 100 | 110 | 120 | 130 | 140 |
| --- | --- | --- | --- | --- | --- | --- | --- | --- | --- | --- | --- | --- | --- |
| Maize | 2015 | A | 0 | 0 | 0 | 0 | 0 | 1 | 1 | 1 | 1 | 2 | 2 |
|  |  | B | 0 | 0 | 1 | 1 | 2 | 2 | 3 | 3 | 3 | 3 | 4 |
|  |  | C | 0 | 0 | 1 | 1 | 2 | 3 | 3 | 3 | 3 | 4 | 4 |
|  | 2016 | A | 0 | 0 | 0 | 0 | 0 | 1 | 1 | 1 | 1 | 2 | 2 |
|  |  | B | 0 | 0 | 1 | 1 | 2 | 2 | 3 | 3 | 3 | 3 | 4 |
|  |  | C | 0 | 0 | 1 | 1 | 2 | 3 | 3 | 3 | 3 | 4 | 4 |
|  | 2017 | A | 0 | 0 | 0 | 0 | 1 | 1 | 1 | 1 | 2 | 2 | 2 |
|  |  | B | 0 | 0 | 1 | 1 | 2 | 2 | 3 | 3 | 3 | 3 | 4 |
|  |  | C | 0 | 1 | 1 | 1 | 2 | 3 | 3 | 3 | 4 | 4 | 4 |
| Cotton | 2015 | A | 0 | 0 | 0 | 0 | 0 | 0 | 1 | 1 | 1 | 1 | 2 |
|  |  | B | 0 | 0 | 1 | 1 | 2 | 2 | 2 | 3 | 3 | 3 | 4 |
|  |  | C | 0 | 0 | 1 | 1 | 2 | 3 | 3 | 3 | 3 | 4 | 4 |
|  | 2016 | A | 0 | 0 | 0 | 0 | 0 | 0 | 1 | 1 | 1 | 2 | 2 |
|  |  | B | 0 | 0 | 1 | 1 | 1 | 2 | 3 | 3 | 3 | 3 | 4 |
|  |  | C | 0 | 0 | 1 | 1 | 2 | 2 | 3 | 3 | 3 | 4 | 4 |
|  | 2017 | A | 0 | 0 | 0 | 0 | 1 | 1 | 1 | 1 | 2 | 2 | 2 |
|  |  | B | 0 | 0 | 1 | 1 | 2 | 2 | 2 | 3 | 3 | 3 | 4 |
|  |  | C | 0 | 1 | 1 | 1 | 2 | 3 | 3 | 3 | 3 | 4 | 4 |

Note: Grade 0 represents intact mulch film with no cracks. Grade 1 represents the appearance of the first crack. Grade 2 represents the appearance of small cracks in 25% of the mulch film in the field. Grade 3 represents the appearance of 2–2.5cm-long cracks. Grade 4 represents the appearance of evenly distributed, network-like cracks. Grade 5 represents that the mulch film has degraded into pieces smaller than 4 cm × 4 cm.

In the maize and cotton treatments, the timing of the appearance of grade 1 cracks in the different mulch films differed each year, which was likely related to factors such as temperature and precipitation in that year. Overall, the common plastic film had grade 1 cracks at 80–100 d, while the biodegradable films had grade 1 cracks at 50–60 d. Therefore, during the early growth stages of maize and cotton, the two biodegradable mulch films had similar effects as the common plastic film in increasing soil temperature and moisture conservation. In addition, during the entire growth period, the plastic film in treatment A only degraded to grade 2 cracks, while in treatment B and C, the mulch films ultimately degraded to grade 4 cracks. In the maize treatment, degradation in treatment C reached grade 3, while treatment B was still at grade 2 at 90 d. In the cotton treatment, in 2015 and 2017, the film degradation at 90 d in treatment C reached grade 3, while in 2016, it was still at grade 2, suggesting that the degradation period of the biodegradable mulch film was not completely consistent in the different crops. For the two crops in most years, the degradation rate was higher in treatment C than in treatment B. This was possibly due to the color of the mulch film, as the black-colored mulch film absorbed more heat itself, which resulted in a relatively faster degradation rate. Within 60 d after the mulch films had been laid, both of the two biodegradable mulch films could retain a relatively intact shape. At 140 d, the two biodegradable mulch films had degraded to grade 4, thus achieving a satisfactory degradation effect in comparison to the common plastic film.

## 4. DISCUSSION

Jie et al.[17], Qiang et al.[18], and Wang et al.[19 found that film mulching could increase crop yield and WUE in potato and maize in semiarid areas in China. However, the application effect of degradable mulch film in arid areas had not been extensively tested. Based on an analysis of the findings of 266 publications, Gao et al.[20] found that the effect of film mulching was significantly better in northern China than in southern China, and the effect was largely dependent on the local climate conditions and management measures. The temperatures of the area of the present study are relatively low in late April, which is unfavorable for maize and cotton germination. However, we found that film mulching drastically increased the crop germination rate, and during the entire growth period, mulch film effectively reduced the time required for crop development. However, as time progressed, the effect of the biodegradable mulch film gradually diminished, which was possibly a result of the high level of degradation at the later stages. In the entire growth period, the plant height, stalk diameter, and leaf area of maize in the different treatments were higher than those in the bare soil, with no significant differences detected among the three films. In cotton, the plant height, leaf area, boll number per plant, and boll weight in the different film mulch treatments were also higher than those in the bare soil, with no significant differences observed among the film types. From the data above, we concluded that the positive effect of biodegradable mulch film on crop plant height, stalk diameter, and leaf area in this region was comparable to that of the common plastic film.

In this study, film mulching significantly increased crop biomass, yield, and WUE. Although there were some differences among the three mulch films in terms of crop yield, these differences were not significant. Due to the stable properties of the common plastic film, it was associated with the greatest increase in crop yield. The black biodegradable mulch film had the second greatest effect due to the longer growth period, while the effects of the clear biodegradable mulch film were slightly smaller than that of the black biodegradable film. Thus, from the long-term perspective of resolving white pollution and reducing the impacts on the environment, these two biodegradable mulch films both have high application value in this region. The increase in WUE from the mulch films was 5% higher in maize than in cotton, which suggests that the applicability of film mulching in maize is slightly better than that in cotton in this region. This might be due to the nature of the plant itself. Xinjiang is a typical arid region, and thus water scarcity is the most important factor limiting the development of local agriculture. Mulch film increased the WUE by 70.35%. In 2017, the agricultural water consumption in this region was 51.440 billion cubic meters (http://data.stats.gov.cn/index.htm). Not considering other losses, the use of mulch films could save approximately 36.2 billion cubic meters of agricultural water usage. This would have great significance for the sustainable development of local agriculture and the protection of the ecological environment.

Due to the limitations in production materials and preparation technologies, biodegradable mulch films often degrade too rapidly, thus having inconsistent effects on crop yield[21,22]. Thus, before biodegradable mulch films can be promoted on a large scale, it is necessary that field experiments are conducted to understand the degradation characteristics of mulch films and whether they are applicable under the local settings. Over a three-year period, we found that both clear and black biodegradable mulch film could retain a relatively intact shape within 60 d after being laid down in the field. Furthermore, their effect on increasing crop yield and WUE was very similar to that of the common plastic film. The replacement of common plastic film with biodegradable mulch films is thus very much applicable in this region.

Based on the agricultural characteristics of this region, the amount of mulch film used in one hectare of soil is 25 kg. When the residual mulch film exceeds 240 kg hm^−2^, it can significantly affect yield[20]. This implies that after nine years of using plastic film, it could negatively impact on crop yield. In addition to the effect of plastic film on crop yield, it can also degrade and produce microplastics[23] which are likely to be transferred into the human body through the food cycle[24], resulting in potential health risks. In this study, after 140 d, the biodegradable mulch films exhibited a significantly higher level of degradation than the plastic film. The ultimate degradation products of the biodegradable film were H_2_O and CO_2_.

Through a three-year field experiment, our study showed that in the short-term, the cultivation of maize and cotton in an arid region using biodegradable mulch film could increase the crop yield and WUE to a similar level as that when using the common plastic film. Considering the environmental benefits and the negative effect of residual film on crop yield, we concluded that biodegradable mulch film was favorable to common plastic mulch film. The effect of biodegradable mulch film under different management conditions was not explored in this study, which is an aspect that could be explored in future research.

## 5. CONCLUSIONS

Through field experiments, we found that film mulching in an arid region was extremely beneficial for crop growth, with the biodegradable mulch films having a similar effect as the common plastic film in the short-term. The three mulch films could significantly increase plant height, stalk diameter, leaf area, crop biomass, and crop yield, and the effects of the biodegradable mulch films were comparable to that of the common plastic film. Additionally, film mulching could significantly increase the WUE in the crops grown in this region, with similar effects observed between the biodegradable mulch film and common plastic film. The degradation properties of the biodegradable mulch films greatly exceeded those of the common plastic film. In addition to increasing crop yield, the biodegradable mulch films are also environmentally friendly. Thus, from a long-term sustainability perspective, the benefits of the biodegradable film outweigh those of the common plastic film.

## Acknowledgments

This research was financially joint supported by the Impact Assessment of Biodegradable Plastic Film on Soil Environment funded by Science&Technology Department of Xinjiang Uygur Autonomous Region under Grant number 2016B02017-4 and West Light Foundation of The Chinese Academy of Sciences under Grant number 2017-XBQNXZ-B-014.

